# Phylogenomics including new sequence data of phytoplankton-infecting chytrids reveals multiple independent lifestyle transitions across the phylum

**DOI:** 10.1101/2023.06.28.546836

**Authors:** Pauline C. Thomé, Justyna Wolinska, Silke Van Den Wyngaert, Albert Reñé, Doris Ilicic, Ramsy Agha, Hans-Peter Grossart, Esther Garcés, Michael T. Monaghan, Jürgen F. H. Strassert

## Abstract

Parasitism is the most common lifestyle on Earth and has emerged many times independently across the eukaryotic tree of life. It is frequently found among chytrids (Chytridiomycota), which are early-branching unicellular fungi that feed osmotrophically via rhizoids as saprotrophs or parasites. Chytrids are abundant in most aquatic and terrestrial environments and fulfil important ecosystem functions. As parasites, they can have significant impacts on host populations. They cause global amphibian declines and influence the Earth’s carbon cycle by terminating algal blooms. To date, the evolution of parasitism within the chytrid phylum remains unclear due to the low phylogenetic resolution of rRNA genes for the early diversification of fungi, and because few parasitic lineages have been cultured and genomic data for parasites is scarce. Here, we combine transcriptomics, culture-independent single-cell genomics and a phylogenomic approach to overcome these limitations. We newly sequenced 29 parasitic taxa and combined these with existing data to provide a robust backbone topology for the diversification of Chytridiomycota. Our analyses reveal multiple independent lifestyle transitions between parasitism and saprotrophy among chytrids, multiple host shifts by parasites, and suggest that the chytrid last common ancestor was a parasite of phytoplankton.

## 1. Introduction

Parasitism is the most successful lifestyle and has emerged many times independently from free-living ancestors (Rueckert et al., 2019). Within major clades, parasitism is often found interspersed with free-living taxa rather than as a lineage-defining trait (Poulin and Randhawa, 2015). Among Chytridiomycota (chytrids), which are early-branching zoosporic fungi that feed osmotrophically via rhizoids, both saprotrophs and obligate parasites are common (distinguished by whether or not they depend on a living host), as are facultative parasites that are able to switch between these lifestyles (Frenken et al., 2017; Powell and Letcher, 2014). Chytrids disperse as free-living motile zoospores that propagate with a single posterior flagellum until they attach to substrate, encyst and grow enzyme-secreting rhizoids into it. The zoospores then develop into sporangia, in which new zoospores are formed by repeated nuclear divisions and eventually released (Medina and Buchler, 2020).

Chytrids fulfil important ecosystem functions. They metabolise many organic compounds, including cellulose, pectin, sporopollenin, keratin and chitin (Ajello, 1948; Chang et al., 2015; Lange et al., 2019; Letcher et al., 2015; Stanier, 1942). As saprotrophs, they decompose recalcitrant organic matter such as insect exuviae, crustacean exoskeletons and pollen (Davis et al., 2019; Powell et al., 2019; Wurzbacher et al., 2014). As parasites, they infect a broad range of hosts comprising plants, amphibians, crustaceans, insects, fungi, phytoplankton and bacteria (e.g. Agha et al., 2016; Dogma and Sparrow, 1969; Longcore et al., 1999; Strassert et al., 2021; van de Vossenberg et al., 2019; Van den Wyngaert et al., 2018; note, due to the lethal effect on their hosts, parasitic chytrids are sometimes also classified as parasitoids). Chytrids thrive in diverse environments such as marine and freshwater aquatic systems and soil (Powell and Letcher, 2014), and occur in climates ranging from the tropics to the Arctic (Hassett and Gradinger, 2016; Jerônimo et al., 2019). As food source, they make difficult-to-access nutrients and energy available for other organisms, thereby influencing zooplankton growth, survival and abundance as part of the so-called mycoloop (Agha et al., 2016; Frenken et al., 2020; Kagami et al., 2014, 2011). Chytrid parasites constitute a substantial proportion of aquatic parasite diversity (Beng et al., 2021; Comeau et al., 2016) and can have significant impacts on biogeochemical cycles and host populations. Phytoplankton-infecting parasites stabilise ecosystems by terminating algal blooms (Frenken et al., 2017; Gleason et al., 2015) and the pathogen *Batrachochytrium dendrobatidis* is a major driver of global amphibian declines (Castro Monzon et al., 2020; McMahon et al., 2013). Other chytrid parasites are of economic importance, causing the wart disease of potatoes (*Synchytrium endobioticum*; van de Vossenberg et al., 2019) or infecting green algae in commercial carotenoid production (*Quaeritorhiza haematococci*; Longcore et al., 2020).

Despite their importance, only a small proportion of chytrid diversity has been described and cultured (Grossart et al., 2019) or has genomic data available for phylogenetic understanding. The simple chytrid morphology limits microscopic assignments of taxonomy, exacerbated by substrate-dependent and intraspecific morphological variation (Hasija and Miller, 1971; Letcher et al., 2006; Paterson, 1963). Attempts to taxonomically and phylogenetically affiliate chytrid species repeatedly reveal that morphology, habitat, lifestyle and substrate or host identity can be misleading traits. This is because convergent adaptations appear to be common (Jerônimo et al., 2019; Powell et al., 2018; Simmons et al., 2021; Van den Wyngaert et al., 2022) and substrate- or host-specificity can vary within taxa depending on season or environment (Frenken et al., 2017; Grossart et al., 2016; Hajek et al., 2013). In most cases, unambiguous taxonomic affiliations should therefore include sequence data (Simmons and Longcore, 2012; Voigt et al., 2021). However, phylogenies based on rRNA genes or a few protein-coding genes have failed to untangle the diversification of chytrids with certainty, leaving the evolution of parasitism within the phylum unresolved. Surprisingly, parasites, such as the amphibian-infecting *B. dendrobatidis*, often cluster with saprotrophs in those phylogenies (James et al., 2006; Joneson et al., 2011; Longcore et al., 2016; Van den Wyngaert et al., 2022). As fungal parasites contribute to major biodiversity losses during the current sixth mass extinction (conspicuous for example within the amphibians; Fisher et al., 2009) and play an important role in a changing world (King et al., 2023), it is crucial to understand the evolutionary dynamics and adaptive potential underlying their emergence.

In this study, we test whether evolutionary transitions from saprotrophy to parasitism have occurred repeatedly within Chytridiomycota by newly sequencing the genomes of 29 parasitic taxa, combining these with existing data from other chytrid saprotrophs and parasites, and using this data to infer a robust phylogenomic tree that allows a comprehensive understanding of chytrid evolution. Based on the number and phylogenetic position of lifestyle transitions uncovered we discuss potential implications for the evolutionary origin of parasitism not only within chytrids but within the fungal kingdom in general.

## 2. Methods

### 2.1 Data collection and generation

We generated new chytrid parasite transcriptomes from laboratory cultures and new genomes from environmental samples. For the transcriptomes, eleven co-cultures of chytrid parasites and their phytoplankton hosts (Appendix A, Table A1) were filtered at the time of highest infection prevalence and sporangia size (filter pore size 0.7 μm, 100–250 mL depending on cell density), and total RNA was extracted using the Transcriptome RNeasy PowerWater Kit (Qiagen, Hilden, Germany) following the manufacturer’s instructions. Libraries were prepared using either the TruSeq stranded total RNA protocol with Ribo-Zero Plus rRNA depletion (*Dangeardia mamillata, Rhizophydium megarrhizum* and *Rhizophydiales* sp. RBA5) or the TruSeq stranded mRNA library protocol (poly A selection; all other cultured chytrids). Transcriptomes were then sequenced (PE 150 bp) at Macrogen Europe (Amsterdam, Netherlands) on the Illumina NovaSeq platform.

For the genomes, phytoplankton from freshly collected lake water samples was screened for chytrid infections. If possible, the chytrid sporangium was detached from the host cell and isolated using two micromanipulators (MMO-202ND; Narishige), each equipped with a microinjector (CellTramm Oil; Eppendorf) holding a capillary (Fig. 1 and Appendix A, Fig. A1). In other cases, chytrids were isolated together with their host cell if the parasite/host ratio was reasonably high enough to expect a sufficient chytrid DNA yield. Genomic DNA was amplified with the REPLI-g Advanced DNA Single Cell Kit (Qiagen), and successful amplification of chytrid DNA was verified by PCR with the primers ITS4ngsF (5’-GCATATCAATAAGCGSAGGA-3’) and LF402R (5’-TTCMCTTTNMRCAATTTCAC-3’; modified from Tedersoo et al., 2015), which target the D1 region of the LSU rRNA gene. Two different template concentrations (1:10 and 1:100) and thermocycler settings were applied to amplify the LSU sequence: One PCR protocol included a denaturation step at 95 °C for 2 min, 35 cycles of 95 °C for 30 s, 53 °C for 30 s and 72 °C for 45 s, and elongation at 72 °C for 5 min. The other protocol included a first PCR of 96 °C for 3 min, 20 cycles of 96 °C for 30 s, 50 °C for 30 s and 72 °C for 60 s, and elongation at 72 °C for 3 min, and a second PCR (using the first product as template), in which the annealing temperature was 55 °C, and the final elongation was 5 min. LSU PCR products were purified and Sanger-sequenced at LGC, Biosearch Technologies (Berlin, Germany), using the same PCR primers (above).

**Figure 1:**
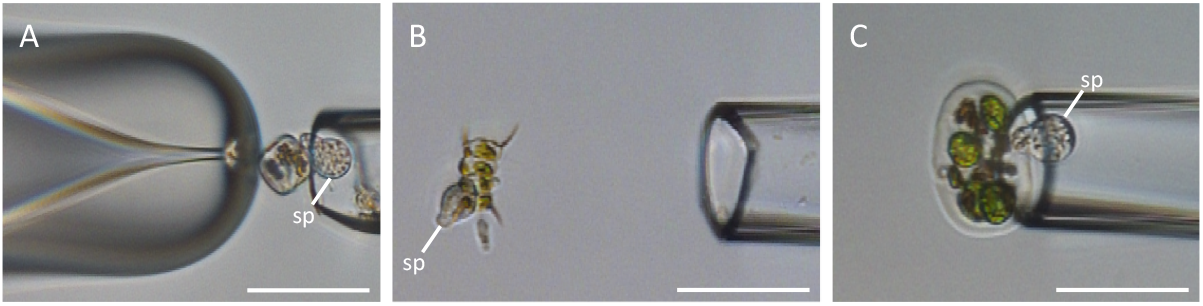
The isolation of chytrid sporangia from environmental samples. Microcapillaries of different shapes and diameters were used to facilitated the detachment of a sporangium (sp) from its host cell. When a sporangium could not be detached from its host, host and parasite were collected together. A – centric diatom, B – *Scenedesmus* sp., C – *Eudorina* sp. Scale bars: 40 μm. The full record for all environmental samples is supplied in Appendix A, Fig. A1.

The LSU sequences were scrutinised by BLASTn searches (Altschul et al., 1990) together with a Maximum Likelihood (ML) tree inference (not shown) using IQ-TREE v. 1.6.12 (Nguyen et al., 2015). A subset of samples was then selected to be whole-genome sequenced in order to represent a broad diversity, with a few exceptions for which two samples with identical LSU sequences were kept in order to increase the genomic data for a certain LSU phylotype. A total of 36 samples were selected for whole genome sequencing. Library preparation and sequencing (PE 150 bp) on the Illumina NovaSeq platform were carried out at Novogene Company Limited (Cambridge, UK).

The High-Performance Computing infrastructure at ZEDAT, Freie Universität Berlin (Bennett et al., 2020), was used for bioinformatic analyses. Reads were trimmed with Trimmomatic v. 0.39 (Bolger et al., 2014) using the options LEADING: 3, TRAILING: 3, SLIDINGWINDOW: 4:15, and MINLEN: 36. Transcriptomes were assembled with Trinity v. 2.10.0 and proteins were predicted with TransDecoder v. 5.5.0 (Haas et al., 2013) using default settings, respectively. Prior to assembly, genomic reads were merged with PEAR v. 0.9.11. (Zhang et al., 2014) and paired reads that remained unmerged were quality filtered with Sickle v. 1.33 with default settings. Genomes were then assembled with SPAdes v. 3.15.5 (Prjibelski et al., 2020) using the single cell option. BUSCO v. 5.1.2 with the database fungi_odb10 (Simão et al., 2015) was used to predict proteins. To determine the host species, morphological features were investigated and BLASTn searches against the SILVA SSU database (Quast et al., 2013) and diamond BLASTx searches (option --very-sensitive) against the NCBI nr database were run using an e-value of 1e-30 each (Buchfink et al., 2021). Similar sequences were clustered using CD-HIT v. 4.8.1 (Fu et al., 2012) with an identity threshold of 85%.

### 2.2 Phylogenomic dataset construction

Our new data was combined with two existing datasets comprising 299 proteins for the phylogenomic analysis. One dataset (Strassert et al., 2021) contained 733 eukaryotic taxa representing all known eukaryote supergroups as well as some prokaryotes and was included to facilitate the identification of contaminants and host sequences from our new data. The other dataset (Strassert and Monaghan, 2022) contained 637 fungal taxa, including 93 chytrids and representatives of other early-branching Holomycota. The broad spectrum of fungal genomes allowed the identification of non-chytrid fungal sequences and provided a framework for quality control and alignment of the new data. Additionally, 66 genomes and transcriptomes of chytrids and other early-branching Holomycota that were publicly available as of August 2022 and were not yet included in the 299-protein datasets were selected from EnsemblFungi, MycoCosm and NCBI, as well as from recent publications (Galindo et al., 2019, 2022; Torruella et al., 2015; Appendix A, Table A2). Raw read data was assembled as described above, with exception that the isolate option in SPAdes was employed in order to assemble downloaded genomic reads. Information about the isolation source of the chosen strains was recorded along with the sequence data: Lifestyle was classified either as saprotrophic, parasitic, i.e. the chytrid overcomes the host’s defence (including facultative parasitism), or as putatively parasitic (e.g. parasites isolated from moribund hosts that possibly lost their defence capability); and habitat was classified as freshwater (isolated from a water sample of a lake, pond or river), soil (including sediment, mud and bogs) or marine.

The two 299-protein datasets were combined, redundant taxa were removed and the resulting dataset served as query to retrieve homologs from the new data using BLASTp searches (BLAST v. 2.7.1, e-value: 1e-20). In a first cleaning round, host sequences, contaminants and deep branching paralogs were removed, while in a second cleaning round, which included only holomycotan sequences and the outgroup, non-chytrid fungi sequences and more recent paralogs were removed by manual inspection of single-protein ML trees. In the first round, sequences were aligned using the -auto function implemented in MAFFT v. 7.475 (Katoh and Standley, 2013) and filtered with trimAl v. 1.4.1 (Capella-Gutiérrez et al., 2009) applying a gap threshold of 0.8. ML trees were inferred with FastTreeMP v. 2.1.11 with the options -lg -gamma -spr 4 -mlacc 2 -slownni (Price et al., 2009). In the second round, which included fewer taxa, computationally demanding but more sophisticated aligning and tree inference methods were used as follows. Sequences were filtered with PREQUAL v. 1.02 (Whelan et al., 2018) using a threshold of 0.95. For global pairwise aligning, MAFFT G–INS–I was then used with a variable scoring matrix (Katoh and Standley, 2016) employing the options --allowshift --unalignlevel 0.6 --maxiterate 0. The alignments were filtered with Divvier v. 1.01 (Ali et al., 2019), using the -partial flag in order to remove further non-homologous residues. After a final trimming step with trimAl (gap threshold: 0.05), single-protein ML trees were inferred with IQ-TREE using best-fitting models according to the Bayesian Information Criterion (BIC), and node support was inferred by ultrafast bootstrap approximation (Hoang et al., 2018) with 1,000 replicates.

The resulting dataset comprised 299 proteins and 765 taxa (including 22 outgroup taxa). The proteins were once more aligned and filtered as described for the second cleaning round and partial sequences belonging to the same taxon that did not show evidence for paralogy or contamination were merged. Proteins were then concatenated into a single matrix (128,829 amino acid positions) with ScaFoS v. 4.42 (Roure et al., 2007). At this step, 10 of the 36 newly genome-sequenced taxa were removed because more than 95% of data was missing. The obtained matrix was used to infer a preliminary tree of the full dataset (Appendix A, Fig. A2) using the site-homogenous model LG+F+G and ultrafast bootstrap approximation (1,000 replicates). Based on this tree, a taxon-reduced dataset was built by merging taxa into operational taxonomic units (OTUs) and by selecting taxa with the most complete sequences to allow for further computationally demanding analyses. Corresponding sequences were newly aligned, filtered, and concatenated as described above (Appendix B). The reduced dataset comprised 107 taxa and 126,130 amino acid positions for the final matrix, including all Chytridiomycota, representatives of the other fungal phyla and of the outgroup (Appendix B). 25 new genomes were merged into 18 taxa by combining virtually identical sequences, and one genome was removed because it did not belong to the chytrids.

### 2.3 Tree inference

Different tree inference methods were applied to the final reduced dataset to uncover potential biases regarding tree topology or statistical support possibly introduced by inference method or evolutionary model. A coalescence tree was computed using ASTRAL-III v. 5.7.7. (Zhang et al., 2018) from 299 single-protein trees that were inferred as in the second cleaning round and partial sequences were merged (see above; Appendix B). An ML tree was computed using IQ-TREE with the best-fitting (according to BIC) heterogenous mixture models LG+C60+G+F (excluding free rate models) and LG+C60+F+R9 (including free rate models) and the posterior mean site frequencies (PMSF) approach (Wang et al., 2018) with 100 bootstrap replicates. A Bayesian Inference (BI) tree was computed using PhyloBayes-MPI v. 1.8 (Lartillot et al., 2013) and the CAT+GTR+G4 model and -dc option. For BI, three independent Markov Chain Monte Carlo (MCMC) chains were run for 2,000 generations and the evolution of the loglikelihood at each sampled point was monitored. The generations before stabilisation were removed as burnin (1,000 generations). The consensus tree did not show global convergence (maxdiff = 1 and meandiff = 0.0157978).

### 2.4 Statistical hypotheses testing

Since ML and BI trees showed conflicts in topology, several statistical tests were carried out to determine the origin of the differences. Incongruent tree topologies were evaluated with the approximately unbiased test (AU-test) in IQ-TREE using the options -n 0 -zb 1,000 -au (Shimodaira, 2002). To distinguish the influence of inference method from that of evolutionary model, another Bayesian tree was reconstructed with PhyloBayes-MPI under the LG+C60+G+F model by running three chains for 3,500 generations and building a consensus tree after a burnin of 2,500 generations (maxdiff = 1, meandiff = 0.0157978). The two evolutionary models, CAT and C60, were compared regarding their fit to the data by posterior predictive analysis in PhyloBayes-MPI employing the -allppred option. Moreover, evolutionary rates for all sites of the matrix were estimated using IQ-TREE with the PMSF tree as a fixed topology and the -wsr option. The fastest-evolving sites were then gradually removed in increments of 10,000 sites and for each stripped alignment, a new ML tree was inferred using IQ-TREE under the LG+C60+G+F model with ultrafast bootstrap approximation (-bb 1,000 -bnni -wbtl). To validate the robustness of the GTR+CAT model to compositional heterogeneity, 25% and 50% of the compositionally most heterogenous sites were removed and two further Bayesian trees were computed from the stripped alignments under the GTR+CAT model. For each tree, three chains were run and a consensus tree was built (25% removed: 2,250 generations, burnin 1,250, maxdiff = 1, meandiff = 0.0157978; 50% removed: 3,000 generations, burnin 1,500, maxdiff = 1, meandiff = 0.0194355).

## 3. Results

We generated new transcriptomes/genomes for 29 chytrid parasites representing seven orders across the entire chytrid phylum (six, in case Sample 34-35 will be assigned to the Lobulomycetales), including four orders for which no such data had been available before: Zygophlyctidales, Polyphagales, Clade I and Sample 34-35. About two thirds of the transcriptomes and genomes showed a high level of completeness (>50%) among the proteins used for our phylogenomic analyses (Appendix A, Table A3). By combining our new data with all chytrid genomes and transcriptomes that were publicly available, we comprehensively represent for the first time chytrid parasite diversity and their evolution with a phylogenomic approach (Figs. 2 and 3).

**Figure 2:**
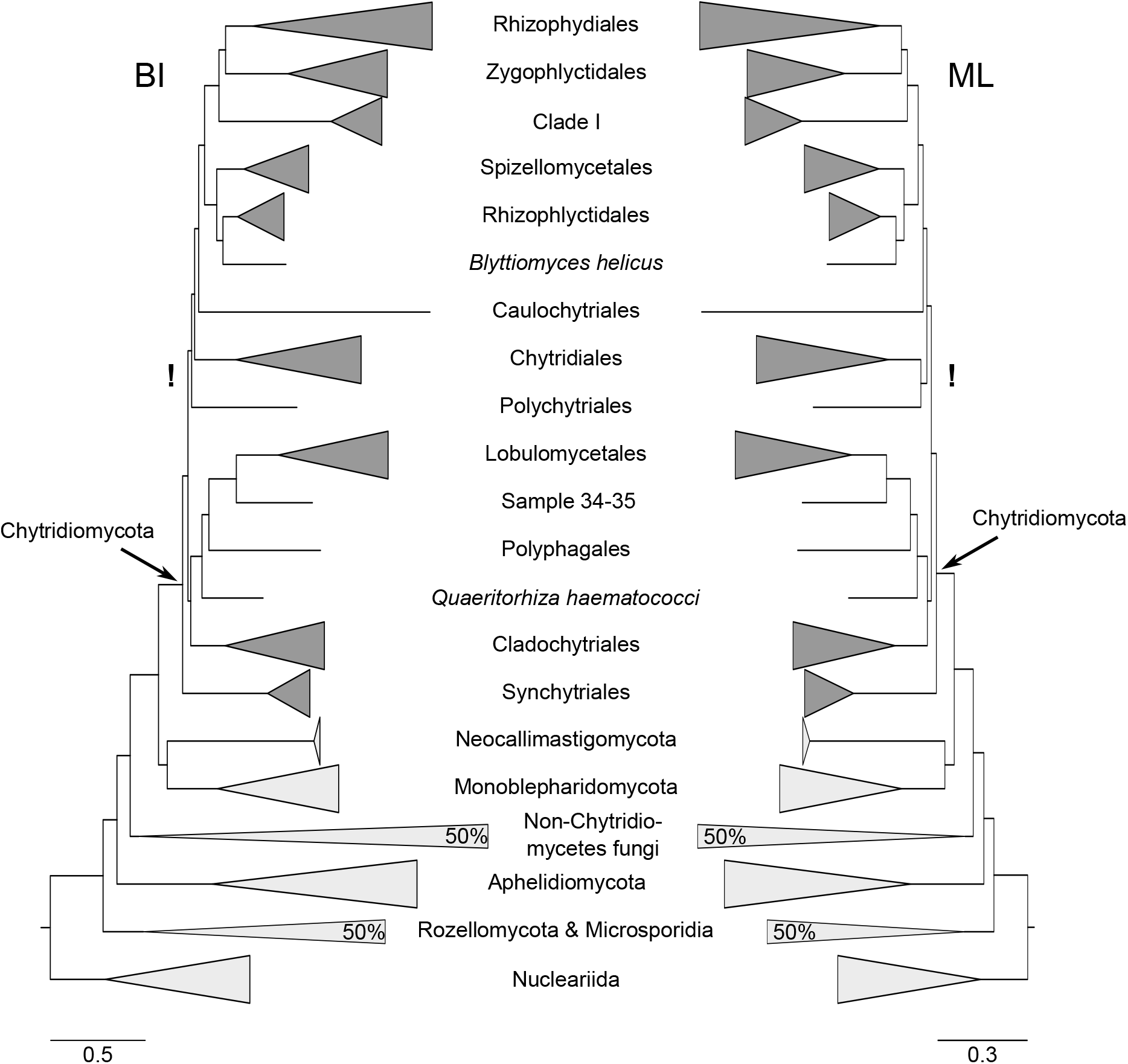
Comparison of tree topologies on the level of orders or phyla between the BI tree (CAT+GTR+G4 model) on the left and the ML tree (LG+C60+G+F-PMSF model) on the right. Trees were inferred from an amino acid alignment of 126,130 positions for 107 taxa. All nodes in both trees were fully supported (posterior probability = 1.0; ML bootstrap = 100). The conflicting placement of Polychytriales either as sister to Chytridiales (ML) or as sister to Chytridiales and its sister clade (BI) is indicated with an exclamation mark. Taxa not affiliated to an order are included with their species binomial. For branches that were reduced in length, reduction is denoted in %. Scale bars indicate amino acid substitutions per site.

**Figure 3:**
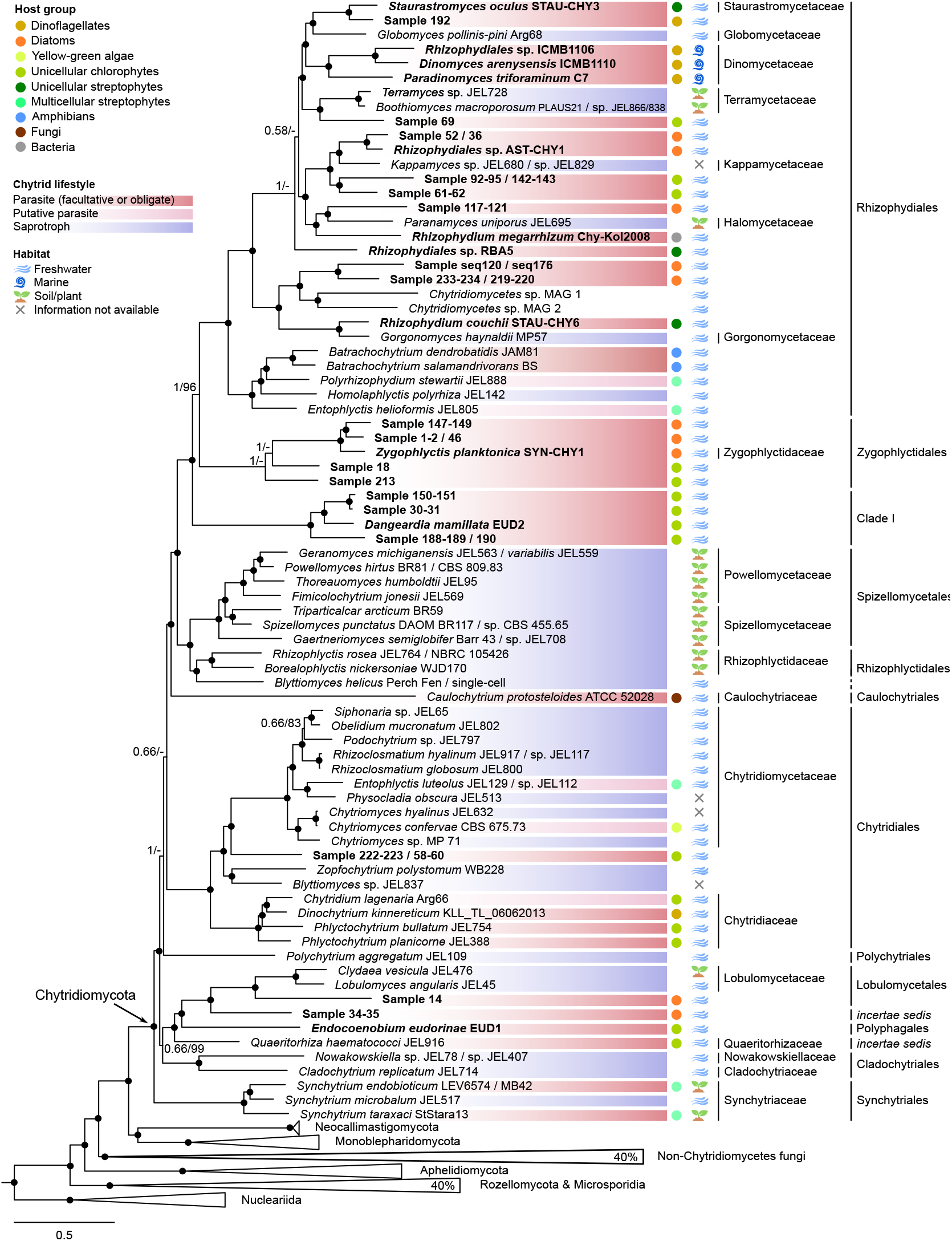
Bayesian tree inferred under the CAT+GTR+G4 model from a matrix of 126,130 amino acid positions and 107 taxa. Newly sequenced taxa are indicated in bold; for cultured taxa, the strain or culture ID is indicated. Nodes that were fully supported both here and in the ML tree (Appendix A, Fig. A3) are indicated by black circles. Nodes that did not receive full support in both trees are denoted with Bayesian posterior probability and ML bootstrap support. For branches that were reduced in length, reduction is denoted in %. Scale bars indicate amino acid substitutions per site. Lifestyle, host association and habitat are depicted for the chytrid strains presented here.

### 3.1 Topology and tree evaluation

Two trees were constructed under site-heterogenous mixture models: One using BI with the CAT+GTR+G4 model (hereafter CATGTR; Fig. 3), and one using ML under the LG+C60+G+F model with posterior mean site frequency profiles (hereafter ML-C60; Appendix A, Fig. A3). The latter was validated with the slightly better-fitting site heterogenous free rate model LG+C60+F+R9 (according to BIC), which showed the same topology but was not used in subsequent analyses due to its high demand for computational resources. Additionally, a multi-species coalescence tree was inferred (hereafter MSC; Appendix A, Fig. A4). The three trees had similar topologies regarding orders and families and most nodes had maximum statistical support (Figs. 2 and 3, Appendix A, Figs. A3 and A4). One conflict between the CATGTR and the ML-C60 trees was the placement of the order Polychytriales, which was represented by only one sequence. While it was sister to Chytridiales in ML-C60 (referred to as Poly-ML topology hereafter; Appendix A, Fig. A3), it branched basal to Chytridiales and its sister clade in CATGTR (referred to as Poly-BI topology hereafter; Fig. 3), each with maximum statistical support. The MSC tree was consistent with the CATGTR tree regarding this topology, although with weak support (Appendix A, Fig. A4). The AU-test rejected the Poly-BI topology but not the alternative (Poly-ML: pAU = 0.999; Poly-BI: pAU = 0.00138). A second conflict concerned one of the new transcriptomes, assigned as *Rhizophydiales* sp. RBA5 (hereafter RBA5), which branched more basal in CATGTR than in ML-C60 and MSC (Fig. 3 and Appendix A, Figs. A3 and A4). Neither topology was rejected by the AU-test (RBA5-ML: pAU = 0.844; RBA5-BI: pAU = 0.156). To test whether the incongruences result from different tree inference methods (ML or BI) or evolutionary model choice (C60 or CAT), another tree using BI under the C60 model was computed, which was congruent to the ML-C60 tree regarding the nodes in question (Appendix A, Fig. A5). Therefore, we compared, which evolutionary model better describes compositional heterogeneity by posterior predictive analysis, which revealed a better fit of the CAT model (Appendix A, Table A4).

To scrutinise the effect of rate heterogeneity on the performance of the C60 model, we progressively removed the fastest-evolving sites from the matrix and computed additional ML-C60 trees from the reduced alignments. This did not lower the bootstrap support for the Poly-ML topology, but rapidly decreased the bootstrap support for the RBA5-ML topology, while the support for the RBA5-BI topology became maximal after the removal of only 30,000 sites (Fig. 4). The CATGTR trees (Appendix A, Figs. A6 and A7), which were reconstructed from alignments of which 25% and 50% of the sites that contribute the most to branch heterogeneity were removed, were congruent to each other regarding the nodes in question and showed robustness of the method to compositional heterogeneity regarding the position of Polychytriales, but not of RBA5, which reflected the same branching as in ML-C60. Biases induced by fast-evolving and compositionally heterogenous sites are therefore unlikely to impact the branching of the single Polychytriales sequence but are most likely sources for the alternative branching of RBA5 (Fig. 4 and Appendix A, Figs. A6 and A7).

**Figure 4:**
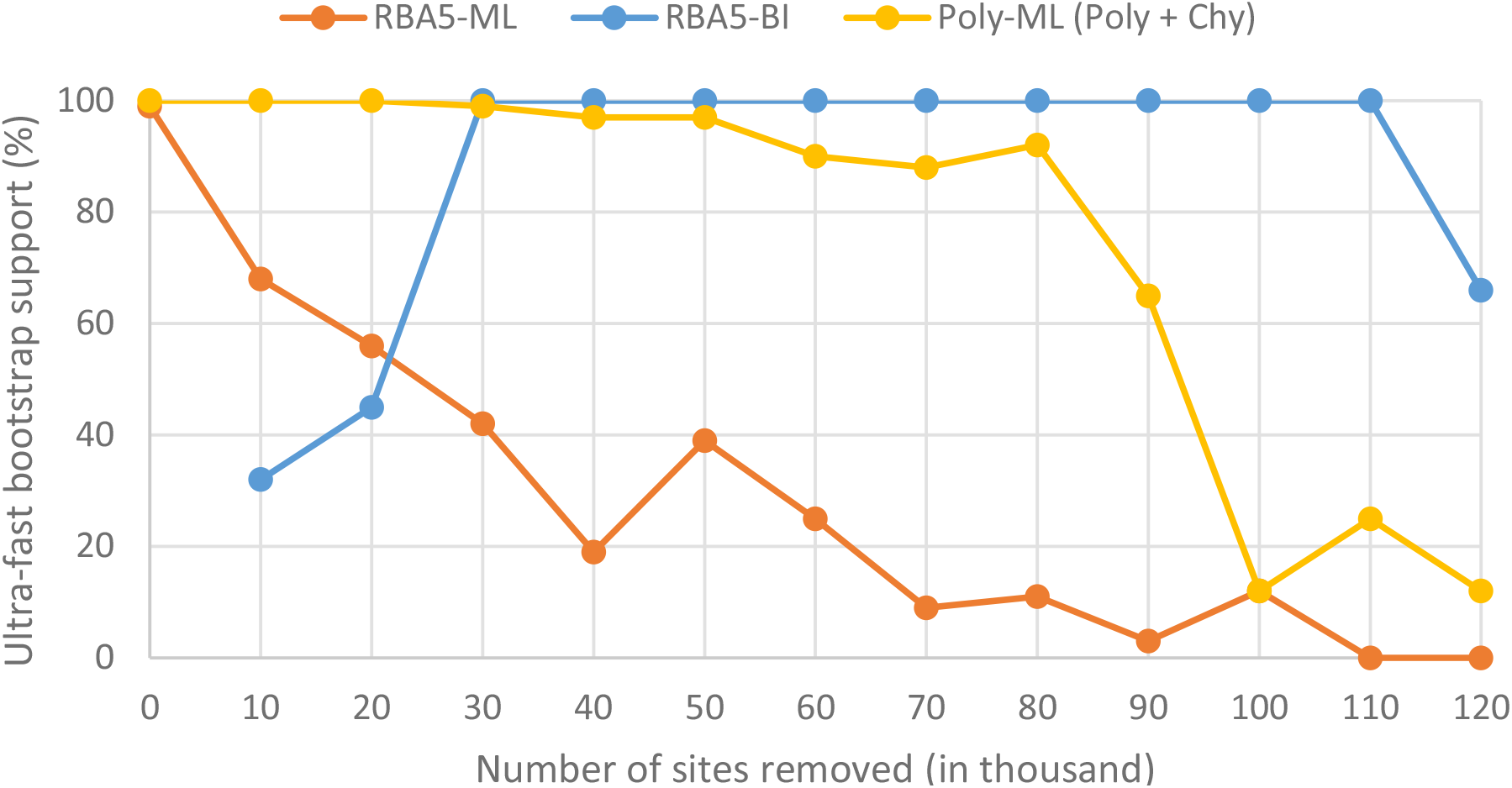
Statistical support for selected tree topologies after removal of the fastest-evolving sites from the alignment in increments of 10,000 sites (of 126,130 amino acid positions in the alignment). Ultrafast bootstrap support is based on 1,000 replicates during ML tree inferences (LG+C60+G+F model).

### 3.2 Chytrid taxonomy

Taxonomic affiliation of previously identified species was based on NCBI taxonomy and previous publications (Galindo et al., 2019; James et al., 2006; Van den Wyngaert et al., 2022). *Incertae sedis* species and our new genomes and transcriptomes were assigned to orders according to their phylogenetic affiliation in the tree (Fig. 3) and Sample 14 was assigned to Lobulomycetales by a BLAST search of the D1 region of the LSU rRNA gene against the NCBI nt data base (uncultured Lobulomycetales; percent identity: 100%; query coverage: 99%). The same search could not determine the order affiliation of Sample 34-35, and neither could a phylogenetic analysis using an LSU dataset, which includes orders of Chytridiomycota that are not represented in our phylogenomic tree (data not shown). The phylogenetic order-level novelty of both *Quaeritorhiza haematococci*, which was basal to Polyphagales, and of Clade I, which was sister to Zygophlyctidales + Rhizophydiales, was verified with maximum support in all analyses, resolving their current *incertae sedis* status proposed by Longcore et al. (2020) and Van den Wyngaert et al. (2022), respectively (Fig. 3). The previously debated position of the order Caulochytriales (Doweld, 2014; Strassert and Monaghan, 2022; Wijayawardene, 2020) was consistent across all analyses, being nested within the Chytridiomycota (as shown by Strassert and Monaghan, 2022) as sister to a clade including Rhizophydiales, Rhizophlyctidales and others.

The debated polyphyletic nature of *Blyttiomyces* was also recovered in our analyses (Blackwell et al., 2011; Powell and Letcher, 2014) and the maximally supported affiliation of *Blyttiomyces helicus* to Rhizophlyctidales resolved its uncertain position for the first time with confidence (Fig. 3), after both molecular and ultrastructural features did not allow an unambiguous placement of this species in the phylogenetic tree (Amses et al., 2022; James et al., 2006; Karpov et al., 2014; Powell and Letcher, 2014). Moreover, whereas *Zopfochytrium polystomum* has been classified as a member of Chytridiaceae based on zoospore ultrastructure and a rather inconclusive rDNA-based phylogeny (Powell et al., 2018), it is shown here (together with *Blyttiomyces* sp.) to be more closely related to a clade comprising the Chytridomycetaceae and Sample 222-223 / 58-60 (Fig. 3).

### 3.3 Parasitism

The majority of phytoplankton parasites that were newly sequenced in this study belong to the orders Rhizophydiales, Zygophlyctidales and Clade I, but one member each of Chytridiales, Lobulomycetales, Polyphagales and one putative order (discovered here by Sample 34-35) were represented as well. While previously only diatom parasites were known in Zygophlyctidales (Seto et al., 2020; Van den Wyngaert et al., 2022), we discovered two chlorophyte parasites. The lifestyle classification that we applied showed that parasitism is common in many chytrid orders and that parasites were often more closely related to free-living saprotrophs than to other parasites (Fig. 3). For example, parasites are the closest relatives of the saprotrophs *Globomyces pollinis-pini, Kappamyces* sp., *Paranamyces uniporus* and *Gorgonomyces haynaldii*. The apparent absence of directional patterns of lifestyle transitions leads to the conclusion that such transitions arose multiple times independently.

Hosts were identified based on morphological characteristics and BLAST searches (newly generated data) and based on the literature (publicly available sequences; Appendix A, Fig. A1 and Table A1). Our results show that host association does not correspond to the phylogenetic affiliation of the parasites and seems to be particularly variable among Rhizophydiales (Fig. 3). Chytrids parasitising on streptophytes, chlorophytes, diatoms and dinoflagellates were often more closely related across these groups than within them. Moreover, the limnic dinoflagellate parasites *Dinochytrium kinnereticum* and Sample 192 clustered within different orders and distinct from the chytrids infecting marine dinoflagellates (*Dinomyces arenysensis, Paradinomyces triforaminorum* and *Rhizophydiales* sp. ICMB1106). The latter formed a comparatively recently diverging monophyletic clade within Rhizophydiales. Within this order, several freshwater phytoplankton parasites were more closely related to soil saprotrophs (e.g. *Boothiomyces, Paranamyces uniporus* or *Gorgonomyces haynaldii*) than to other freshwater parasites. Among Synchytriales, habitat transition correlated with a change in lifestyle. Adaptations to different environments did therefore not appear related to the chytrids’ phylogeny, indicating their convergent nature.

## 4. Discussion

Parasitism is a common lifestyle among chytrids and is a key reason for their global importance as regulators of phytoplankton populations and as pathogens of plants and animals. To understand the evolution of parasitism within the phylum, we produced new genomes and transcriptomes of 29 phytoplankton-infecting chytrid taxa. These represent seven orders allowing the first comprehensive analysis of chytrid lifestyle evolution to date. The provided backbone topology was constructed under site-heterogenous mixture models and remained robust to exhaustive statistical scrutiny with only one exception on the order-level (Polychytriales). A discrepancy to a previously published multi-protein ML phylogeny was given by the placement of Caulochytriales: Sister to Synchytriales (Amses et al., 2022) *versus* sister to a clade comprising Rhizophydiales, Zygophlyctidalis, Clade I, Spizellomycetales and Rhizophlyctidales (this study). Both the more comprehensive taxon-sampling in this study as well as the application of site-heterogenous mixture models, which usually better fit such heterogenous data, are likely sources for this incongruence. We found that conflicts between the CATGTR and ML-C60 trees were the result of differences in evolutionary models, which is in line with previous studies showing the CAT model to better describe both compositional heterogeneity across branches and rate heterogeneity across sites (e.g. Strassert and Monaghan, 2022). At the same time, the unresolved conflicts also show that describing compositional heterogeneity can be challenging for both models when data is scarce. Nonetheless, even considering the discussed methodological challenges, the conclusions about the evolution of lifestyles across the chytrid phylum as inferred here remain robust.

Our phylogenomic tree revealed a remarkable number of independent lifestyle transitions between saprotrophy and parasitism in chytrids. More independent transitions are likely to be uncovered when sequence data is available for lineages that are not represented here or only represented by members with the same lifestyle: For example, insect-infecting nephridiophagids (associated with Cladochytriales) or parasitic Spizellomycetales species are not represented here, nor are those members of *Kappamyces* and *Terramyces* that were observed in lab experiments to parasitize on diatoms and cyanobacteria, respectively (Letcher et al., 2006; Reñé et al., 2022; Strassert et al., 2021; Wakefield et al., 2010). Also, the parasite *Dinomyces arenysensis* is known to grow on pine pollen as well (Fernández-Valero, personal communication). Furthermore, the number of transitions between lifestyles is underestimated here because facultative parasitism cannot always be distinguished from obligate parasitism and of those species that are found on moribund hosts it cannot conclusively be determined whether they caused its death or responded to it (Alster and Zohary, 2007; Longcore, 1995; Paulitz and Menge, 1984; Simmons et al., 2021). Similarly, for the uncultured parasites sequenced here, we could not determine host fitness at the time of infection, but given the sporangia size and host state at the time of collection, we conclude that parasitic interactions with a living host must have occurred. Facultative parasitism may be an adaptation to a periodic absence of hosts or substrates, or they may be strategies to fill different niches (Frenken et al., 2017; Van den Wyngaert et al., 2022). In particular in extreme environments like the Arctic with seasonal varying nutrient availability, the capability to switch between lifestyles is highly advantageous (Kellogg et al., 2019).

We found chytrid host groups were paraphyletic and occurred within multiple orders, indicating that these adaptations arose several times independently. A wide phylogenetic distribution of chytrids associated with one particular host group has also recently been inferred by rRNA gene phylogenies, albeit with lower support (Fernández-Valero et al., 2022; Van den Wyngaert et al., 2022). Within the dinoflagellate parasites, the only group for which both freshwater and marine members were included in our analysis, adaptation to the same host group was found even across different habitats. It is safe to assume that more independent transitions from both soil and freshwater to marine habitats associated with adaptations to certain host groups will emerge from metabarcoding surveys (as recently shown for dinoflagellate parasites; Fernández-Valero et al., 2022) but also in robust phylogenomic studies when data from more species becomes available. In this context it is also noteworthy that here, we present a more conservative estimate of independent transitions as classifications concerning lifestyle, host association and habitat were applied for the particular strains included in this study, regardless of what is known about other representatives of the same species or genera.

In contrast to the unique evolutionary transition to pathogenicity in the amphibian infecting *Batrachochytrium* species, the frequent occurrences of phytoplankton parasitism that we observed in most chytrid orders interspersed with saprotrophic taxa are unlikely to be based on comparable lingeage-specific extensions of repeat-rich regions and gene family radiations including genes that are not present in saprotrophic relatives (Farrer et al., 2017; Joneson et al., 2011; Wacker et al., 2023). Instead, the necessary functional gene repertoire of the diverse forms of chytrid phytoplankton parasitism is more likely to be of ancient origin predating chytrid diversification and might have been differentially retained in extant lineages. Both free-living saprotrophy and phytoplankton parasitism in chytrids probably rely on ancestral trait sets given the frequency of transitions in both directions.

Parasitism is common across the entire fungal kingdom and occurs in other phyla, such as in Zoopagomycota and Dikarya (Ahrendt et al., 2018; Naranjo-Ortiz and Gabaldón, 2019a and 2019b; Poulin and Randhawa, 2015), in similar patterns as shown here for chytrids. Already before the emergence of fungi, endobiotic phagotrophic predation involving parasitic interactions likely was the ancestral lifestyle during early holomycotan evolution, as inferred from extant parasitic Rozellomycota, Microsporidia and Aphelidiomycota (Galindo et al., 2022; James et al., 2013; Torruella et al., 2018). The same is assumed for a potentially aphelid-like fungal ancestor that lived in association with an ancestral streptophyte lineage (Berbee et al., 2020; Chang et al., 2015; Keeling and Fast, 2002; Torruella et al., 2018). Following these assumptions, the osmotrophic lifestyle evolved inside the protected space of a phytoplankton host cell before the parasitic fungal ancestors became independent of the intracellular stage and eventually of the parasitic association (Galindo et al., 2022; James et al., 2013). As a major evolutionary novelty, osmotrophy allowed fungi to adapt to diverse organic substrates and enabled them to abandon endoparasitism in favour of saprotrophy, thereby breaking Dollo’s law (Cruickshank and Paterson, 2006; Naranjo-Ortiz and Gabaldón, 2019). Based on the wide distribution of parasitism across the chytrid phylum and based on considerations of the fungal ancestor being a parasite of microalgae, we propose that the ability to parasitize phytoplankton is the ancestral state of chytrids that has been lost multiple times independently. The ancestral lineage would have retained its capability to live independently of a host, allowing for subsequent lifestyle transitions to facultative parasitism or saprotrophy, as happened frequently in Rhizophydiales. The Zygophlyctidales, in contrast, are represented only by obligate parasitic lineages with high host specificity, making it likely that members of this clade became highly dependent on their hosts. While ancestral state reconstructions on our topology using an ML-based approach (Ishikawa et al., 2019) were inconclusive (data not shown), a reversal to a free-living state is known to be more common than traditionally assumed (Cruickshank and Paterson, 2006; Xu et al., 2016). The frequency with which host dependence was reversed in chytrids is striking, since parasites generally evolve reduced genomes and lose genes coding for essential functions for which they depend on the host (Poulin and Randhawa, 2015), such as in Microsporidia (James et al., 2013). Potential ancestral chytrid parasites, in contrast, would have retained a genomic repertoire enabling transitions to the free-living lifestyle of extant saprotrophs. We propose this as a hypothesis to be tested in future studies.

## Conclusion

We obtained a broad picture of chytrid parasite diversity and host associations using single-cell genomics on environmental samples, which proved to be a valuable complement to our culturing efforts and transcriptome sequencing. The robust backbone topology for chytrid diversification presented here indicates that phytoplankton parasitism is the ancestral lifestyle of chytrids, while pathogenicity to metazoa is a more recently derived evolutionary novelty. Chytrid ecology is demonstrated to be lineage specific and independent of phylogenetic affiliation, because transitions in lifestyle and host association have occurred often and convergently. Understanding the ecological diversity and adaptive potential within the phylum is important in the light of emerging chytrid pathogens in a changing world.

## Data availability

Ribosomal marker genes have been submitted to NCBI under the accession numbers ###. All other data used in this study can be found in Appendix B.

**Appendix A: Supplementary material:**Supplementary figures and tables

**Appendix B: Supplementary data:**Alignments and single-protein trees

## CRediT authorship contribution statement

**Pauline C. Thomé:** Data curation, Formal analysis, Investigation, Resources, Software, Validation, Visualisation, Writing -original draft, Writing -review & editing.

**Justyna Wolinska:** Conceptualisation, Resources, Writing -review & editing.

**Silke Van Den Wyngaert:** Resources, Writing -review & editing.

**Albert Reñé:** Resources, Writing -review & editing. **Doris Ilicic:** Resources, Writing -review & editing. **Ramsy Agha:** Resources, Writing -review & editing.

**Hans-Peter Grossart:** Resources, Writing -review & editing.

**Esther Garcés:** Resources, Writing -review & editing.

**Michael T. Monaghan:** Conceptualisation, Resources, Writing -review & editing.

**Jürgen F. H. Strassert:** Conceptualisation, Data curation, Funding acquisition, Investigation, Methodology, Project administration, Resources, Software, Supervision, Writing -original draft, Writing -review & editing.

## Acknowledgements

We thank Katrin Preuß for her support in identifying potential chytrid hosts based on morphological features. We are also grateful to Jordina Gordi for her support in maintaining the marine cultures used in this study. We wish to thank Elisabeth Funke, who extracted RNA of three of the here presented samples. J.F.H.S. acknowledges support from the German Research Foundation (DFG; grant STR1349/2-1, project no. 432453260) and the German Federal Ministry of Education and Research (BMBF; Förderkennzeichen 033W034A). AR and EG acknowledge the institutional support from the “Severo Ochoa Centre of Excellence“ accreditation (CEX2019-000928-S).

## Appendix A

### Supplementary figure legends

**Figure A1:** Host–Chytrid pairs from environmental samples that were used to generate sequence data. Host identifications were based on morphological characteristics in combination with BLASTn and diamond BLASTx searches against the SILVA SSU database and the NCBI nr database, respectively. With exception of Sample 192 and Sample 213, which were collected from lake Jungfernheide (Berlin, Germany), all other samples were collected from lake Müggelsee (Berlin, Germany). Scale bars: 50 μm. **A:** Sample 1-2, centric diatom, same chytrid taxon as B. **B:** Sample 46, centric diatom, same chytrid taxon as A. **C:** Sample 14, *Asterionella formosa*. **D:** Sample 30-31, *Chlamydomonas* sp. **E:** Sample 34-35, centric diatom. **F:** Sample 36, centric diatom, same chytrid taxon as G. **G:** Sampe 52, centric diatom, same chytrid taxon as F. **H:** Sample 58-60, *Scenedesmus* sp., same chytrid taxon as I. **I:** Sample 222-223, *Scenedesmus* sp., same chytrid taxon as H. **J:** Sample 61-62, *Scenedesmus* sp. **K:** Sample 92-95, *Scenedesmus* sp. (or *Elakatothrix* sp.), same chytrid taxon as L. **L:** Sample 142-143, *Scenedesmus* sp., same chytrid taxon as K. **M:** Sample 117-121, centric diatom (putative *Melosira* sp.). **N:** Sample 147-149, *Stephanodiscus* sp. **O:** Sample 150-151, *Chlamydomonas* sp. **P:** Sample 188-189, putative *Lobomonas* sp., same chytrid taxon as Q. **Q:** Sample 190, *Lobomonas* sp., same chytrid taxon as P. **R:** Sample 192, *Peridinium* sp. **S:** Sample 213, *Pediastrum* sp. **T:** Sample 219-220, centric diatom, same chytrid taxon as U. **U:** Sample 233-234, centric diatom, same chytrid taxon as T. **V:** Sample seq120, centric diatom, same chytrid taxon as W. **W:** Sample seq176, centric diatom, same chytrid taxon as V. No pictures were taken from Sample 18 (putative *Planktosphaeria* sp.) and Sample 69 (putative *Coelastrum* sp.). For information about the host identities of other parasites represented in this study, please see Table A1.

**Figure A2:** ML tree inferred under the site-homogenous LG+F+G model from the full dataset (765 taxa). Ultrafast bootstrap support is reported from 1,000 replicates. Sequences coloured in silver were removed to build the reduced dataset used for the final trees and sequences coloured in cyan were merged with their neighbours into OTUs. For newly sequenced taxa, provisional working names are given.

**Figure A3:** ML tree inferred under the site-heterogenous LG+C60+G+F-PMSF model from the reduced dataset (107 taxa). Bootstrap support is reported from 100 replicates. For newly sequenced taxa, provisional working names are given.

**Figure A4:** Coalescence tree based on 299 single-protein trees inferred with ML from the reduced dataset (107 taxa) under site-heterogenous models that were best-fitting according to BIC. Node support indicates support per quadripartition without bootstrapping. For newly sequenced taxa, provisional working names are given.

**Figure A5:** BI tree inferred under the site-heterogenous LG+C60+G+F model from the reduced dataset (107 taxa), consensus of three independent runs. Node support values are Bayesian posterior probabilities. For newly sequenced taxa, provisional working names are given.

**Figure A6:** BI tree inferred under the site-heterogenous GTR+CAT model from the reduced dataset (107 taxa) after removing 25% of the compositionally most heterogenous sites, consensus of three independent runs. Node support values are given by Bayesian posterior probabilities. For newly sequenced taxa, provisional working names are given.

**Figure A7:** BI tree inferred under the site-heterogenous GTR+CAT model from the reduced dataset (107 taxa) after removing 50% of the compositionally most heterogenous sites, consensus of three independent runs. Node support values are given by Bayesian posterior probabilities. For newly sequenced taxa, provisional working names are given.

## Supplementary table legends

**Table A1**: Parasitic chytrid species and their hosts. Information is shown for taxa of which the genomes/transcriptomes were publicly available and for cultures from this study. For information about the environmental samples from this study, please see Fig. A1.

**Table A2**: Lineage information and sequence data source of all taxa that were not included in Strassert et al. (2021) and Strassert and Monaghan (2022).

**Table A3**: Average data completeness in the final reduced alignment across all proteins according to ScaFoS.

**Table A4**: Z-scores obtained by posterior predictive analysis with PhyloBayes.

